# Galanin Analogs Prevent Seizure-Induced Respiratory Arrest

**DOI:** 10.1101/2022.03.21.485198

**Authors:** Ryley Collard, Miriam C. Aziz, Kevin Rapp, Connor Cutshall, Evalien Duyvesteyn, Cameron S. Metcalf

## Abstract

**Objective:** Sudden Unexpected Death in Epilepsy (SUDEP) accounts for 20% of mortality in those with recurrent seizures. While risk factors, monitoring systems, and standard practices are in place, the pathophysiology of SUDEP is still not well understood. Better knowledge of SUDEP and its potential mechanisms of action is crucial to reducing risk in this patient population and developing potential treatment options. Clinical studies and animal models of SUDEP suggest that diminished post-ictal respiratory control may be the dominant mechanism contributing to mortality. Recently, it was demonstrated that the depletion of the neuropeptide galanin in the amygdala occurs in human SUDEP. The amygdala plays a key role in the central integration of respiratory signaling; the depletion of galanin may represent a critical change that predisposes individuals to SUDEP.

**Methods:** To evaluate the potential benefit of enhancing galaninergic signaling as a means to protect against SUDEP, we studied seizure-induced respiratory arrest (S-IRA) following central (intracerebroventricular, intra-amygdala) and systemic (intraperitoneal, subcutaneous) administration of galanin agonists. Seizure naïve and seizure experienced (fully kindled) mice were tested.

**Results:** Central and systemically-administered galanin analogs protect against S-IRA in naïve C57Bl/6J mice. Differential efficacy between receptor subtype-selective analogs varied based on the route of administration. Sub-chronic systemic administration at doses that reduced 6 Hz seizures also protected against S-IRA. Acute treatment benefits also extended to fully kindled mice subjected to tonic extension.

**Significance:** These data demonstrate that galanin agonists may be protective against post-ictal respiratory collapse.

**KEY POINTS:**

- Central and systemic galanin agonists prevent seizure-induced respiratory arrest.
- Efficacy was observed in three separate mouse strains under various experimental conditions.
- Sub-chronic administration demonstrated galanin agonist protection against respiratory arrest.
- Acute systemic administration also conferred protection against respiratory arrest following tonic extension.
- Galanin analogs may represent a novel potential therapy in SUDEP-susceptible individuals.

## INTRODUCTION

Epilepsy can lead to a variety of serious complications and is associated with a reduction in life-expectancy. One of the leading causes of mortality in those with recurrent seizures is an event known as Sudden Unexpected Death in Epilepsy (SUDEP)^1-5^. A pivotal clinical study demonstrated that post-seizure (post-ictal) apnea precedes asystole^6^, suggesting that respiratory control may be a major mechanism in SUDEP. Further, clinical observations and mouse models suggest that respiratory neurocircuitry is altered in individuals susceptible to early mortality^7-11^. Death after seizures may result from a combination of poor respiratory responses and failure in normal arousal mechanisms that promote breathing^2, 3, 7, 12-14^.

Generalized tonic-clonic seizures (GTCS) are a major risk factor of SUDEP and may lead to hypoxia, hypercapnia, failed arousal mechanisms, and thus an inability to recover breathing following seizures^2, 3, 15, 16^. A GTCS can spread to other brain regions, including those involved in breathing. Several forebrains and brainstem nuclei may regulate breathing following seizures. One important area of interest is implicated in post-ictal respiration is the amygdala. Apnea results when seizures spread to the amygdala, and amygdala lesions reduce seizure-induced respiratory arrest (S-IRA) in mice^17-19^. Moreover, the amygdala has direct connections with brainstem breathing control sites^20-22^. The amygdala is enriched in neuropeptides that modulate neuronal activity and can be depleted following seizures^23^. Interestingly, depletion of the neuropeptide galanin in the amygdala occurs in human SUDEP^23^. Galanin is an anticonvulsant neuropeptide activated under conditions of enhanced neural activity^24^ and plays a critical role in responses to respiratory stressors^25, 26^. Further, galanin is robustly upregulated after various insults such as nerve injury and seizures^27-29^. Similarly, galanin is increased in the amygdala following seizures^30^. Therefore, the amygdala is a critical relay point between seizures and downstream breathing control following seizures, and the depletion of galanin may represent a critical pathophysiologic change predisposing some individuals to SUDEP. It may also be that enhancement of galanin signaling in the amygdala protects against respiratory collapse following seizures.

There still remains no effective evidence-based treatment or prevention against SUDEP. The mainstay of management has been addressing modifiable risk factors such as ensuring medication adherence, but with few results^2, 3, 13, 31^. Novel therapies may therefore prove beneficial if they offer benefits targeted explicitly to SUDEP risk, in addition to any potential benefits of reducing seizure burden. Therefore, the purpose of this study is to use rodent models to demonstrate how galanin agonists may have neuroprotective effects in preventing mortality in mice experiencing seizure-induced respiratory arrest (S-IRA). Respiratory arrest, arising following tonic extension elicited in mice, has been used as a model of SUDEP and recapitulates apnea following GTCS^32, 33^. Further, this model offers the opportunity to screen compounds with acute and chronic effects at preventing S-IRA. These studies described herein will therefore demonstrate the potential utility of galanin agonists in restoring respiration after seizures in naïve and seizure-experienced mice.

## MATERIALS AND METHODS

### Animals

Three different mouse strains were used: C57BI/6J (5-6 weeks old, Jackson Laboratory, Bar Harbor, ME, USA), CD-1 (5-7 weeks old, Charles River Laboratories, Kingston, NY, USA), and CF-1 (5-7 weeks old, Charles River Lab, Kingston, NY, USA). Animals were allowed free access to food and water, except during testing periods. Prior to testing, animals were allowed 1 week to acclimate to housing conditions. All mice were housed in plastic cages in rooms with controlled humidity, ventilation, and lighting (12 hours on – 12 hours off). The animals were housed and fed in a manner consistent with the recommendations in the “Guide for Care and Use of Laboratory Animals” (National Research Council). Housing, handling, and testing was performed in accordance with Public Health Service policy guidelines and a protocol approved by the Institutional Animal Care and Use Committee of the University of Utah.

### Compound Preparation/Administration

810-2 (Gal_2_-preffering) and 505-5 (Gal_1_-preferring) (molecular weights 2124 g/mol and 2113 g/mol, respectively)^34-38^ was synthesized by PolyPeptide Laboratories (San Diego, CA USA). 810-2 was dissolved in a vehicle (VEH) solution of 20% hydroxy propyl beta cyclodextrin (HPβCD; Sigma, St. Louis, MO, USA) in acetate buffer [0.1 M acetic acid (from glacial acetic acid stock; Sigma, St. Louis, MO, USA); 0.1 M sodium acetate (Sigma, St. Louis, MO, USA), pH 4.5]. 505-5 was dissolved in a VEH solution of 1% Tween 20 (Sigma, St. Louis, MO, USA) in 0.9% NaCl. Compounds were administered by intraperitoneal (IP), intracerebroventricular (ICV; free-hand injection), or intra-amygdala (1-2 weeks post unilateral amygdala cannulation). Separate experiments were also conducted for administration of galanin analogs by subcutaneous (SC) sub-chronic (14-day) administration. 810-2 and 505-5 were prepared in 50:50 dimethylsulfoxide:water at concentrations of 67.2 mg/ml and 16.8 mg/ml (respectively), and placed in implantable mini-pumps (Alzet, model 1002)

### Surgical Implantation of Unilateral Amygdala Cannulae

Procedures are similar to those described previously for implantation of unilateral cannulae for drug administration^39^. Buprenorphine (0.01-0.2 mg/kg) was administered 1h prior to surgery. Isoflurane (2-5% in O_2_) was used for anesthesia, and mice were placed in a stereotaxic apparatus. A Dremel drill was used to drill a single hole over the right hemisphere followed by placement of a 22-gauge cannula above the dura mater (AP −1.2, ML 3.3). This guide cannula was glued in place using dental acrylic. Antibiotic ointment was applied to the surface around the head cap and penicillin (60000 units, SC) was administered. Mice were then allowed to recover in their home cages (singly housed) for ∼1 week.

### Surgical Implantation of Osmotic Minipumps

Alzet minipumps were prepared and immersed in saline overnight. Under isoflurane anesthesia (2-5% in O_2_), as described above, a small incision was made between the scapulae (shaved prior to incision) followed by clearance of subcutaneous fascia with forceps. Minipumps were placed in the subcutaneous space and the incision sutured with surgical silk. Antibiotic ointment was placed around the incision and mice were allowed to recover. Mice were monitored daily for the duration of the 14-day treatment.

### Acute Electrical Seizure Induction and Post-Stimulation Monitoring

#### Maximal Electroshock Seizure Model

MES has been used as a model of seizure induction for preclinical pharmacology studies routinely for many years^40^. Furthermore, tonic extension following MES stimulation is followed by apnea and death in some mouse strains and has been used as a model of S-IRA^32^. Prior to testing, tetracaine (0.5% in saline) was applied to the corneas. MES seizures were induced using a 50 mA current (0.2 sec duration) via corneal electrode^41, 42^. This stimulation intensity produces tonic extension in nearly all mice tested. Following stimulation, mice were observed for the presence of tonic extension and the resumption of post-ictal breathing. Mice not displaying tonic extension were considered protected.

#### 6 Hz Seizure Induction

This seizure assay has been used as a model of preclinical pharmacology studies in mice^42^ and was used in these studies to confirm efficacy observed following treatment with galanin analogs^38, 43, 44^. Seizures were induced using the 32 mA stimulus intensity (3sec, 6 Hz, 32 mA) via corneal electrodes. Prior to testing, tetracaine (0.5% in saline) was applied to the corneas. Seizure induction is associated with characteristic behaviors including jaw and forelimb clonus with or without loss of righting and rear limb clonus. Animals not displaying any of these behaviors were considered protected.

### Respiratory Monitoring

CF-1 mice subjected to maximal electroshock were also monitored for cardiorespiratory activity before and following seizures using a MouseOx (Starr Life Sciences) monitoring system. On the day prior to testing, mice were anesthetized with 2-5% isoflurane and the neck region shaved with a surgical razor. On the following day, prior to seizure induction, mice were acclimated to pulse oximetry collars for ∼15 min followed by a baseline recording session (5 min). A plastic sham collar (not connected to monitoring hardware) was placed initially, followed by oximetry collars and baseline recording sessions. After the baseline recording, mice were subjected to maximal electroshock stimulation followed by a post-stimulation recording session (5 min). Oximetry data were extracted as text files and transferred to Microsoft Excel databases for data analysis. A 30-second recording epoch was selected from each recording session and heart rate, respiratory rate, and oxygen saturation (SpO_2_) values averaged over this period.

### Intraperitoneal Administration

810-2 and NAX 505-5 were administered to C57BI/6J and CD-1 mice 1h prior to MES stimulation This time point was selected based on previous studies demonstrating peak efficacy for these analogs in the mouse 6 Hz assay 1h following IP administration^38, 44^. 810-2 was administered at doses of 8, and 16 mg/kg IP (N= 10-24 per group), doses were selected based on previous *in vivo* efficacy studies for this compound^36, 45^. 505-5 was administered at doses 2, and 4 mg/kg IP (N= 9-24 per group). Doses for each compound were selected based on previous *in vivo* efficacy studies^36-38, 45^. A separate group of mice were treated with VEH (N=21-24), IP administration, 1h pre-treatment time.

### Central Administration

Mice received a single administration centrally of either 810-2, 505-5, or VEH. A 5 μl injection volume was used for all central injection studies, and mice only received one injection before being tested and euthanized. Doses of 1-4 nmol were used for intracerebroventricular (ICV) administration whereas a larger dose range was used for intra-amygdala (IA) treatment. For ICV administration a 10 μl Hamilton syringe with a 25-gauge needle was used for administration. Mice were placed under gentle restraint and a free-hand injection was made through the skull surface and into the lateral ventricular space. By contrast, for IA injection studies, mice were implanted 1-2 weeks prior with cannulae centered unilaterally over the amygdala and an injection needle was used to administer compounds over a 30sec period. Initial low doses of 810-2 were selected (0.02-2 nmol) to evaluate tolerability to injection of peptides in this brain region. Following a maximal dose (4.7 nmol) was used, comparable to that used for ICV studies. Treatments occurred 15 min prior to testing for all central injections.

### Statistical Analysis

The data was presented as means ± standard error. A comparison between two means was performed using a Student’s t-test, and multiple comparisons were made using a one-way ANOVA followed by a Newman-Keuls or a Dunnett’s test for a posteriori analysis of the difference between group means. A Fisher’s exact test was used to indicate appropriate sample sizes. Survival analyses were analyzed using a log-rank (Mantel-Cox) test. Results where a P < 0.05 was considered significant.

## RESULTS

### Galanin analogs reduce seizure-induced respiratory arrest in mice: differential effects of route of administration and analog

Galanin analogs 810-2 and 505-5 were administered to C57Bl/6J mice using different routes of administration, followed by MES stimulation for evaluation of S-IRA. First, each analog was administered 1h prior to testing using IP administration: 810-2 8,16 mg/kg, 505-5 (2, 4 mg/kg) vs VEH. All mice tested demonstrated a characteristic tonic extension following MES^40, 46, 47^. Typically, breathing is absent or diminished during tonic extension in all mice, and many mice die as a result, particularly for CD-1 and C57Bl/6J strains. As shown in Table 1, VEH treatment was associated with a high mortality rate (62%, 23/37 mice died). Systemic administration of 810-2 decreased mortality at a dose of 16 mg/kg (25%, 5/20 died; P<0.05, Fisher’s exact test) whereas 505-5 was without significant effect at the doses tested. As higher doses of each analog are associated with untoward effects (sedation, lethargy), additional doses were not included. CD-1 mice respond similarly to testing, and both 505-5 and 810-2 significantly improve mortality in these mice (505-5, 4 mg/kg IP, P<0.05 vs VEH, Fisher’s exact test; 810-2, 16 mg/kg, P<0.05 vs VEH, Fisher’s exact test; Table S1).

**Table 1.** Systemic and central administration of galanin analogs: effect on (S-IRA) in C57B1/6J mice.

| Compound | Systemic (IP) | <u>Mortality (# died/N)</u> |  |
| --- | --- | --- | --- |
|  |  | Central (ICV) | Central (IA) |
| VEH | 23/37 | 13/16 | 11/17 |
| 810-2 | 8mg/kg: 11/19 | 1 nmol: 6/11 | 0.02 nmol: 6/7 |
|  | 16mg/kg: 5/20* | 2 nmol: (10/16) | 0.1 nmol: 8/8 |
|  |  | 4 nmol (11/16) | 0.2 nmol: 9/9 |
|  |  |  | 4.7 nmol: 6/23* |
| 505-5 | 2 mg/kg: 9/20 | 1 nmol: 11/16 | 4.7 nmol: 9/21 |
|  | 4 mg/kg: 8/20 | 2 nmol: 4/8 |  |
|  |  | 4 nmol: 4/11* |  |

\*P<0.05, \*\*P<0.01, Fisher's exact test vs. VEH. Routes of administration: IP (intraperitoneal), ICV (intracerebroventricular), IA (intra-amygdala).

Following systemic administration studies, ICV administration was performed. Previously, doses of 1-2 nmol (ICV) demonstrated efficacy against 6 Hz seizures for galanin analogs^44^. Therefore, doses in this range were used for these studies. Similar to that observed following systemic administration, ICV VEH was associated with a high mortality rate (81%, 13/16). While the 810-2 was without significant effect using the doses tested, 505-5 reduced seizures at the highest dose tested (4 nmol; 4/11, *P<0.05 vs VEH, Fisher’s exact test). Next, using unilateral intra-amygdala cannulae, galanin analogs were administered by direct injection in freely-behaving mice. Initially, a dose of VEH was administered and showed a high mortality rate (65%; 11/17). Following several low doses of 810-2 were administered (0.02-0.2 nmol) to determine whether direct injection would produce untoward effects. As these concentrations were well-tolerated, a higher concentration (4.7 nmol) was administered for both 505-5 and 810-2. 810-2 significantly reduced mortality (26%, 6/23; P<0.05 vs VEH, Fisher’s exact test), whereas 505-5 was not significantly effective (43%; 9/21).

### The Gal_2_-preferring analog 810-2 prevents mortality following sub-chronic subcutaneous administration

We sought to determine whether sub-chronic (14 days) systemic administration would produce similar efficacy against S-IRA. These galanin analogs have a short half-life (1-2h), thereby requiring repeated (multiple times/day) injections, implantable minipumps were used to provide a more consistent compound exposure. Therefore, SC minipumps were prepared to administer daily doses of 4 or 16 mg/kg of 505-5 and 810-2, respectively. Galanin receptors are G-protein coupled transmembrane proteins and develop tolerance to repeated agonist exposure warranted verification of efficacy^24, 48^. Because these analogs are effective in the mouse 6 Hz seizure model^38, 43, 44^, a single stimulus was administered at the end of the study on the day prior to MES testing. As shown in Table 2 for the 6 Hz assay, 505-5 and 810-2 both improved seizure outcomes, with 505-5 producing superior efficacy (21/26 protected, P<0.001 vs VEH), but 810-2 also proving efficacious (6/11 protected, P<0.05 vs VEH). On the following day, MES was performed it was observed that a majority of VEH-treated mice died following tonic extension (75%, 12/16). Also, while 505-5 did not improve mortality (69% mortality, 18/26), 810-2 significantly reduced S-IRA (18%, 2/11, P<0.01 vs VEH) (Table 2).

**Table 2.** Reduced S-IRA following sub-chronic administration of galanin analogs in C57Bl/6J mice.

| **Groups** | **Dose (mg/kg)** | **6 Hz Efficacy**<br>**(# protected/N)** | **% Mortality**<br>**(# died/N)** |
| --- | --- | --- | --- |
| VEH | 0 | 2/16 | 75 (12/16) |
| 505-5 | 4 | 21/26*** | 69 (18/26) |
| 810-2 | 16 | 6/11* | 18 (2/11)** |

\*P<0.05, \*\*P<0.01, \*\*\*P<0.001 vs VEH, Fisher's exact test.

### Evaluation of post-ictal hypoxia, respiration, and heart rate following MES seizures

In order to study effects on postictal respiration following MES, we evaluated SpO2, respiratory rate, and heart rate in CF-1 mice. Notably, this strain of mice survives TE despite having a period of apnea following seizures. VEH-treated CF-1 mice subjected to MES-induced tonic extension experience a brief period (10-20sec) of apnea and decreased SpO2, which coincides with hyperventilation (see Figure 1). There were no major changes in heart rate observed.

**Figure 1.**
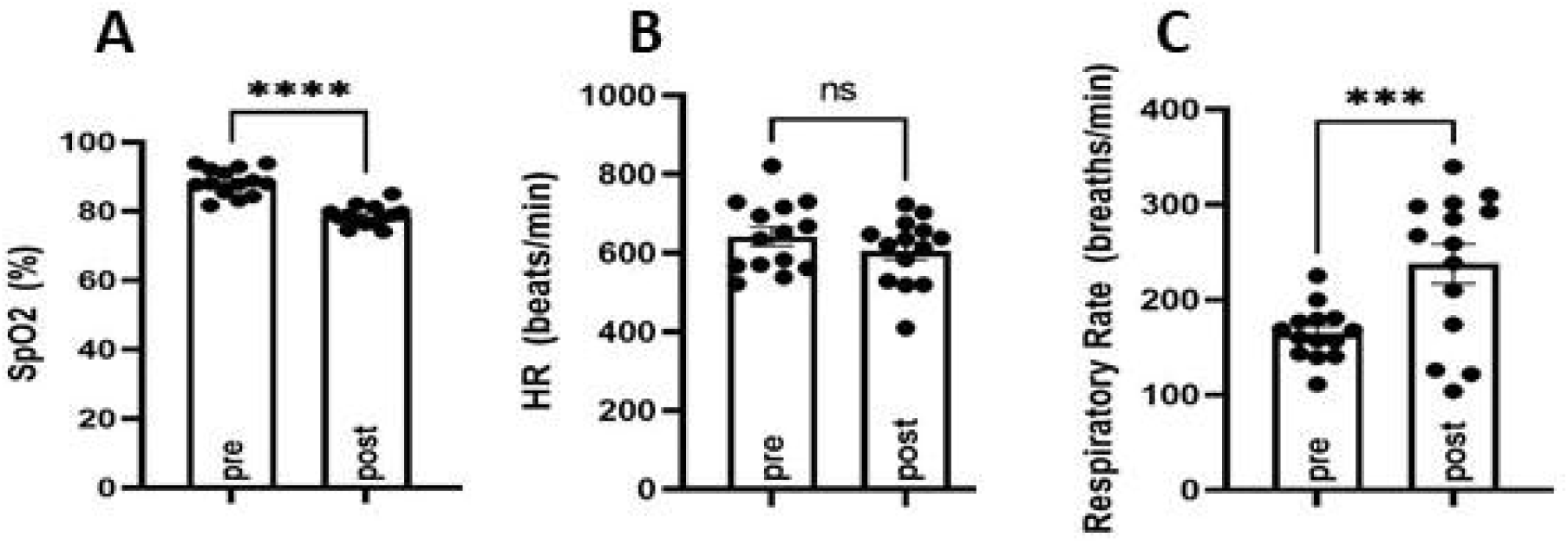
Evaluation of oxygen saturation (SpO2), respiratory rate, and heart rate before (pre) and following (post) MES-induced tonic extension in CF-1 mice. SpO2 was diminished (A) concomitant with an increased respiratory rate (B) and no major changes in heart rate (C). ***P<0.001, ****P<0.0001 compared to pre, Student’s t test. N=14.

Following a baseline MouseOx analysis, galanin analogs were administered by IP injection 1h prior to testing. VEH, 810-2 (dose range 4-16 mg/kg), and 505-5 (dose range 1-4 mg/kg), were administered prior to a single MES stimulation and 5 min observation using MouseOx. 810-2 was without major effect on post-ictal hypoxia (SpO2), hyperventilation (respiratory rate), or heart rate (Fig 2A-C). By contrast, 505-5 worsened hypoxia at 2 mg/kg (Fig 2D) but was without major effect on respiratory rate or heart rate (Figs 2E-F).

**Figure 2.**
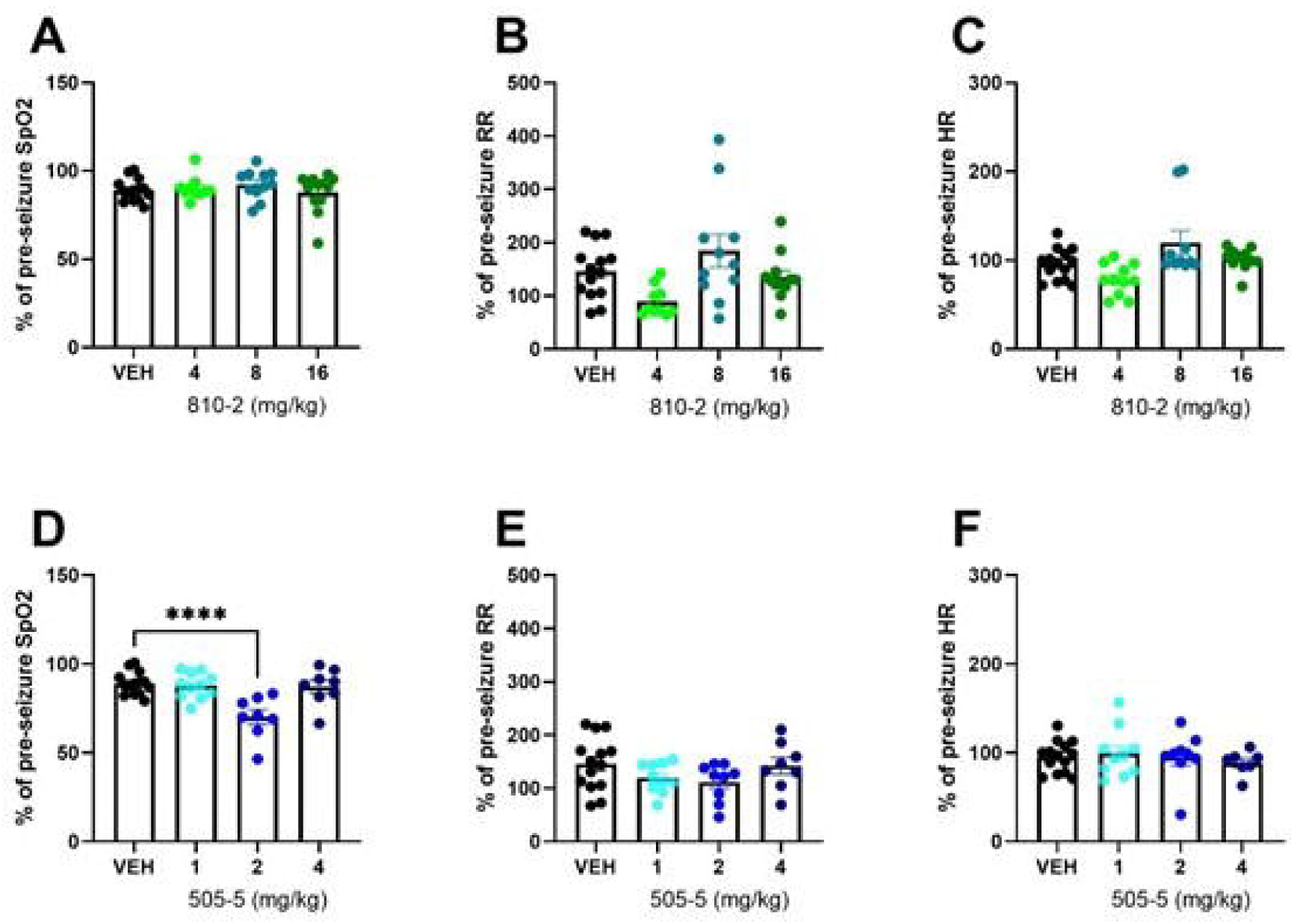
Evaluation of cardiorespiratory parameters following tonic extension in CF-1 mice. Data are presented as a percentage of baseline (pre-seizure) values. VEH-treated values are the same as shown in Figure 1, except expressed as a percent of baseline. 810-2 (A-C) and 505-5 (D-F) were administered by IP injection 1h prior to tonic extension and testing (MouseOx). Oxygenation (SpO2; A, D), respiratory rate (RR; B, E), and heart rate (HR; C, F) were assessed for all animals. N=8-14. ****P<0.0001 (One-way ANOVA, Sidak’s multiple comparison post-hoc test).

### Galanin analogs reduce seizure-induced respiratory arrest in fully kindled mice

#### Evaluation of corneal kindling as a potential model of SUDEP

We obtained kindling records from the NIH Epilepsy Therapy Screening Program for several cohorts of CF-1 mice subjected to the corneal kindling paradigm. Data presented here are from naïve kindled mice not treated with any investigational compounds. We observed that fully kindled mice have a high mortality rate, particularly when tonic extension occurs after aquiring full kindled status (Figure S1). In separate studies, a cohort of CF-1 mice were kindled and subjected to baseline and post-ictal (MES assay) cardiorespiratory evaluation (MouseOx). We also observed that although SpO2 was similarly reduced in age-matched control (CONT) and fully kindled (KIND) mice following TE, respiration did not significantly increase in KIND mice, though this change was present in mice that were partially kindled (PART; i.e. received daily stimulations but failed to acquire full kindled status) (Figure S2). Interestingly, heart rate was increased in KIND mice following TE, whereas it was unchanged in other groups.

#### Evaluation of the effect of galanin analogs on S-IRA in fully kindled mice

As a comparator to studies in naïve C57Bl/6J mice, a cohort of mice from this strain was fully kindled and subjected to treatment with galanin analogs using MES-induced TE. Initially, a group of untreated kindled mice was subjected to TE and it was observed that a majority (6/9 animals died following TE). Similarly, VEH-treated kindled mice also demonstrate a high mortality after TE (11/16 died) (Table 3). 505-5 improved mortality following TE in kindled mice (4/17 died) (Table 3). While 810-2 had a similar magnitude of effect, the difference was not significant compared to VEH (Table 3).

**Table 3.**
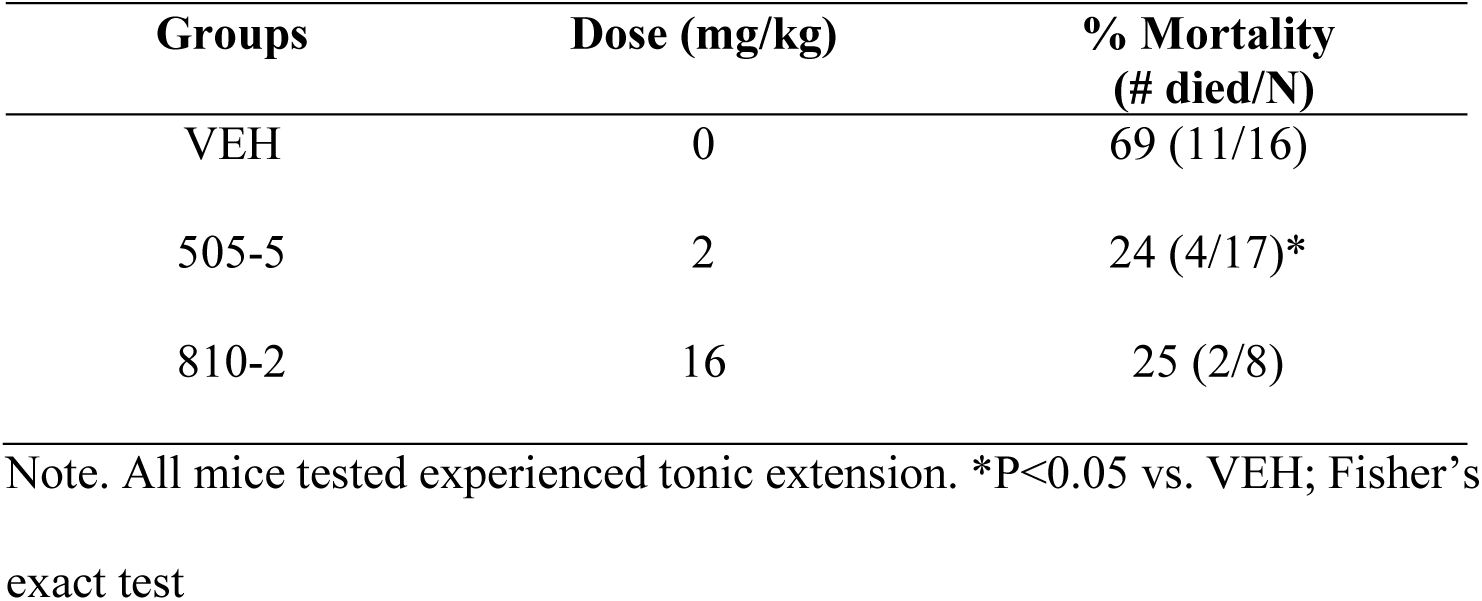
Reduced S-IRA following a single administration of galanin analogs in fully kindled C57Bl/6J mice.

## DISCUSSION

SUDEP is a major cause of mortality in epilepsy, and better understanding of mechanisms of death may aid in the development of novel therapies to prevent or reduce the likelihood of death following seizures. Recent studies in human SUDEP have suggested a potential role for the neuropeptide galanin in the amygdala. Therefore, we sought to further evaluate the role of galanin in a mouse model of S-IRA under various conditions. Our principal findings include (1) galanin analogs reduced S-IRA in naïve mice following systemic and central administration, (2) 810-2 prevented mortality following sub-chronic administration, (3) protective effects of systemic galanin analogs were confirmed in fully kindled mice.

Poor adherence to treatment or pharmacoresistance may increase the occurrence of GTCS in patients with various forms of epilepsy. Further, GTCS are a critical risk factor for SUDEP. While genetic and acquired epilepsy models may recapitulate seizures and mortality in epilepsy, drug screening in these models may be challenging due to the need for prolonged administration and monitoring to determine whether treatment reduces seizure burden, tonic extension, and mortality. This kind of study may be critical in demonstrating translational relevance for novel therapies, but throughput may be low. However, acute treatment and testing models may be more amenable to drug screening and help identify lead candidates for more rigorous testing. Therefore, we applied our previous knowledge of galanin analogs to a S-IRA model to better understand potential benefits of galanin-based therapies on reducing the likelihood or onset of SUDEP. Systemic administration of receptor subtype preferring analogs suggested that targeting of Gal_2_ may be beneficial, as 810-2 demonstrated superior reduction in S-IRA over 505-5. Of note, the systemic doses used for each compound were not raised beyond those used in the study, as higher doses are associated with untoward effects (sedation and motor impairment, data not shown). Therefore, future endeavors with galanin analogs may seek to optimize administration through combinatorial pharmacology. For example, recent work has clearly demonstrated a potential benefit for serotonergic therapies in reducing the incidence of SUDEP in animal models^32, 33, 49, 50^. While beyond the scope of this study, combined treatment with serotonin-and galanin-based therapies may yield an added benefit.

We sought to extend our systemic administration studies by using central injection as a means to more specifically target regions where galanin may exert protective effects against S-IRA. Galanin is expressed throughout the forebrain neurocircuitry, including several sites that affect cardiorespiratory and autonomic function. ICV administration studies suggest an important role for Gal_1_ over Gal_2_ in reducing S-IRA. Conversely, Gal_2_ played a more prominent role when galanin analogs were administered directly into the amygdala. The amygdala has emerged as an important regulator center for the interaction of seizures with respiratory control. Apnea results when seizures spread to the amygdala and amygdala lesions reduce seizure-induced respiratory arrest (S-IRA) in mice^17-19^. Additionally, the amygdala has reciprocal connections with brainstem respiratory centers^20-22^. Importantly, galanin is depleted in the amygdala in human SUDEP^23^, which suggests that supplemental galanin in this region may reduce the incidence of SUDEP. The reason behind these discrepancies in our ICV and IA data are unclear but may involve multiple neural circuits affecting respiration after seizures. Furthermore, it is important to recognize that IA administration was centered over the basolateral amygdala but may have also spread to the central amygdala. The central amygdala may play an important role in the spread of a seizure via the amygdala in suppressing respiration. Future work may include application of galanin analogs in a more subregion-specific and receptor subtype-specific manner.

Tonic extension is lethal in a variety of mouse strains. We evaluated tonic extension following MES stimulation in C57Bl/6J, CD-1, and CF-1 mice. Both C57Bl/6J and CD-1 mice experience similar mortality following MES stimulation; we observed different responses to galanin compounds. Where 810-2 (Gal_2_-preferring) was effective in both strains in reducing S-IRA. 505-5 (Gal_1_-preferring) was only effective in CD-1 mice. The mechanism for this strain-dependent discrepancy is unknown and may be likely to different expression in brainstem respiratory centers between the two species. Future studies may further explore these potential differences. Conversely, CF-1 mice generally survive tonic extension. Interestingly, we observed that this mouse strain demonstrates respiratory distress (diminished SpO_2_, hyperventilation) following tonic extension. Thus, tonic extension is a major respiratory stressor, yielding periods of apnea that may be insurmountable for some mouse strains. We hypothesized that galanin compounds may prevent mortality by preventing post-ictal hypoxia and therefore we evaluated oxygenation, heart rate, and respiratory rate in CF-1 mice treated prior to MES stimulation. Neither galanin compound prevented post-ictal hypoxia at any dose tested. Further, 505-5 demonstrated an exacerbation of this response (2 mg/kg). Therefore, the protective effects of galanin compounds in S-IRA may not be due to a direct effect on preventing the cardiorespiratory response to apnea.

G-protein coupled receptor may undergo internalization following stimulation with agonists^51^. In order to explore the use of galanin compounds as potential therapeutic agents, we performed sub-chronic administration using implanted minipumps pre-filled with either 810-2 or 505-5. To confirm antiseizure efficacy at the doses selected, we first performed 6 Hz seizure testing and observed that both 810-2 and 505-5 retained efficacy in this assay. On the following day, mortality following tonic extension was assessed, and we observed that only 810-2 was effective in reducing mortality. The effect of 810-2 in this assay is consistent with the acute (single administration, IP) effect of this compound and confirms that systemic administration of Gal_2_-preferring compounds exerts a protective effect against S-IRA.

Corneal kindling has long been used as an assay for studying epileptogenesis, epilepsy pathology, and antiseizure drug pharmacology. We observe that in CF-1 mice, a portion of animals die during and following kindling acquisition. We obtained several historical data cohorts from the ETSP (NIH, NINDS) Contract Site (University of Utah) and reviewed kindling histories for the presence of tonic extension after daily kindling stimulation. We reviewed data from naïve kindled mice and we surmised that while naïve CF-1 mice may be resistant to death following tonic extension, this may shift in a kindled state. We identified several animals that died after having experienced one or more bouts of tonic extension. Deaths observed in this cohort did not necessarily occur immediately following tonic extension. Further, these data are consistent with clinical observations that the presence of GTCS are a major risk factor for SUDEP. To better understand the effect of tonic extension on kindled mice, we subjected kindled CF-1 mice to pulse oximetry recordings before and after MES stimulation. We observed that oxygenation drops following tonic extension in kindled mice in a similar manner to age-matched control mice. Interestingly, hyperventilation observed in control mice was not observed to the same extent in kindled mice, though heart rate was diminished in these animals. Reduced post-ictal hyperventilation among kindled mice may contribute to an increased risk for mortality in some animals.

To extend our studies of galanin analogs as protective of S-IRA in naïve mice, we also evaluated these analogs in fully kindled C57Bl/6J mice. In contrast to observations in naïve mice, where 505-5 was ineffective in reducing S-IRA, 505-5 reduced mortality (2 mg/kg). These data suggest a potential shift in receptor expression in brain respiratory centers toward Gal_1_ sensitivity (e.g. receptor upregulation or downregulation of Gal_2_), though this was not examined in the present study. Future studies will examine the expression of each galanin receptor subtype in kindled mice.

In summary, we have observed that galanin compounds protect against S-IRA following tonic extension in naïve and kindled mice. This was observed after systemic and central administration, and responses varied depending on route, location, strain, seizure history, and receptor subtype preference. While these studies suggest the potential translational benefit for galanin in SUDEP prevention, additional work is needed to differentiate the utility of these analogs in SUDEP-susceptible individuals.

## ACKNOWLEDGEMENTS

The authors would like to thank Dr. Misty Smith (Dept. of Pharmacology &Toxicology, University of Utah), for her assistance with ICV injections of galanin analogs. We thank the Epilepsy Therapy Screening Program of the National Institutes of Neurological Diseases Disorders and Stroke for providing kindling history data in untreated CF-1 mice used as part of these studies.

