## Supplemental for "Galanin Analogs Prevent Seizure-Induced Respiratory Arrest"

**SUPPLEMENTAL MATERIALs**

**Table S1.** Reduced S-IRA following systemic administration of galanin analogs in CD-1 mice.

| Groups | Dose (mg/kg) | Tonic Extension (# observed/N) | % Mortality (# died/N) |
|---|---|---|---|
| VEH | 0 | 24/24 | 71 (17/24) |
| 505-5 | 4 | 24/24 | 38 (9/24)* |
| 810-2 | 16 | 23/23 | 35 (8/23)* |

*P<0.05 vs VEH; Fisher's exact test.

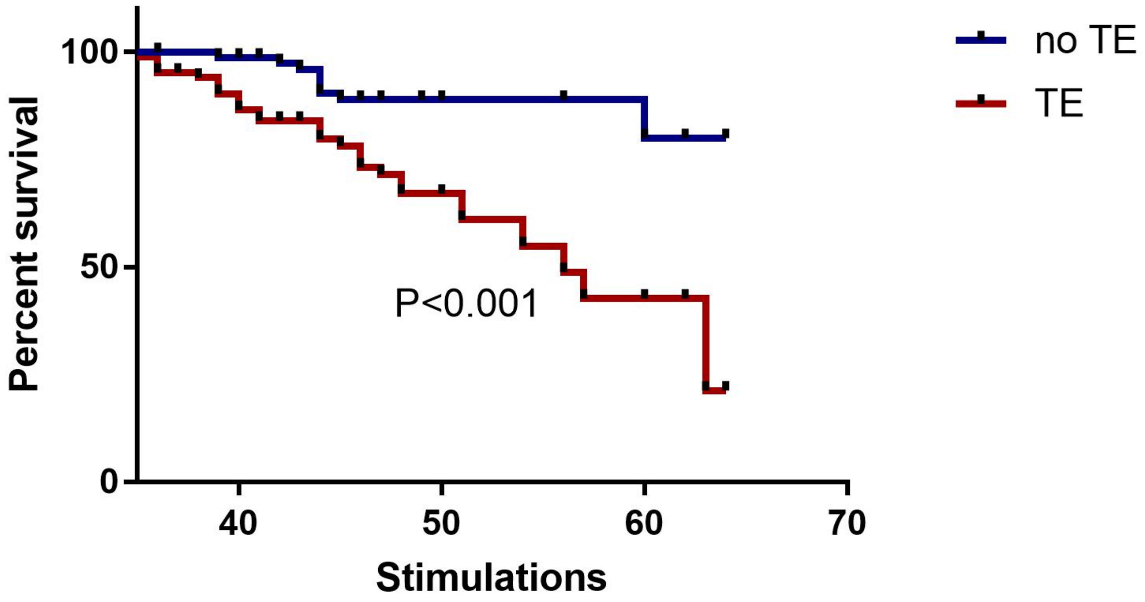


**Figure S1**. Diminished survival observed in fully kindled CF-1 mice that experience tonic extension. A data mining effort was conducted to identify mice that experience at least one tonic extension event following daily kindling stimulations. When plotted as a group of mice with tonic extension (TE) and without (no TE), it was observed that survivability was greatly diminished if tonic extension had occurred. N=425 mice (no TE n=265 (52 died, 213 survived), with TE n=160 (53 died, 107 survived). P<0.001 (Log Mantel-Cox test).

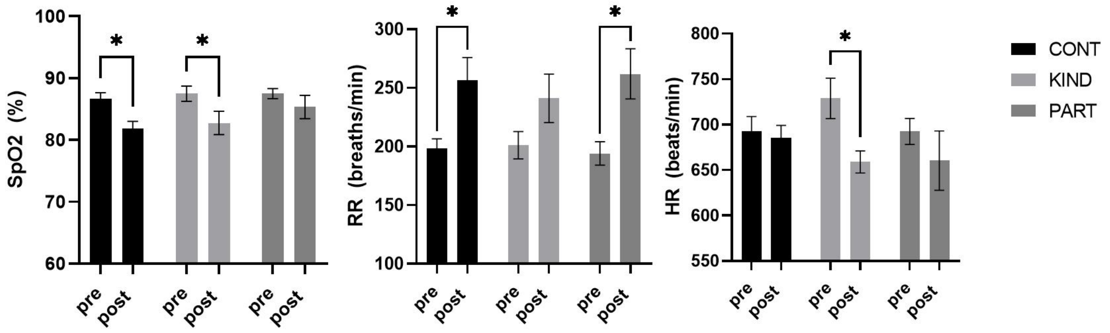


**Figure S2**. Evaluation of oxygen saturation (SpO2), respiratory rate (RR) and heart rate (HR) in fully kindled mice (CF-1). SpO2 (A), RR (B), and HR (C) values were obtained for age-matched control (CONT), fully kindled (KIND), and partially kindled (PART) mice. PART mice received daily kindling stimulation (as for KIND mice) but failed to reach kindling criterion. *$P<0.05$, 2-way ANOVA, Sidak's multiple comparison test. N=7 per group.
